# Antigen mobility regulates the dynamics and precision of antigen capture in the B cell immune synapse

**DOI:** 10.1101/2024.10.31.621246

**Authors:** Hannah C. W. McArthur, Maro Iliopoulou, Anna T. Bajur, Katelyn M. Spillane

## Abstract

B cells discriminate antigens in immune synapses by capturing them from antigen-presenting cells. This discrimination relies on the application of mechanical force to B cell receptor (BCR)-antigen bonds, allowing B cells to selectively disrupt low-affinity interactions while internalizing high-affinity antigens. Using DNA-based tension sensors combined with high-resolution imaging, we demonstrate that the magnitude, location, and timing of forces within the immune synapse are influenced by the fluidity of the antigen-presenting membrane. Transitioning antigens from a high-mobility to a low-mobility substrate significantly increases the probability and speed of antigen extraction while also improving affinity discrimination. This shift in antigen mobility also reshapes the synapse architecture, altering spatial patterns of antigen uptake. Despite these adaptations, B cells maintain consistent levels of proximal and downstream signaling pathway activation regardless of antigen mobility. They also efficiently transport internalized antigens to major histocompatibility complex class II (MHCII)-positive compartments for processing. These results demonstrate that B cells mount effective responses to antigens across diverse physical environments, though the characteristics of that environment may influence the speed and accuracy of B cell adaptation during an immune response.

## Introduction

Protective antibody responses begin when B cells capture antigens from antigen-presenting cells (APCs). Upon binding antigens through their B cell receptors (BCRs), B cells internalize and process these antigens, ultimately displaying antigenic peptides on major histocompatibility complex class II (MHCII) molecules (1, 2). This presentation elicits T cell help, which—combined with BCR-induced metabolic and transcriptional activation—drives B cell expansion and differentiation into plasma, memory, or germinal center cells (3–9). Through class switch recombination and somatic mutation, activated B cells generate diverse BCRs, establishing a pool of cells that produces broad and potent antibodies and maintains immunological memory (10). Without T cell help, however, antigen-stimulated B cells undergo apoptosis (11). Thus, a B cell’s fate hinges on its ability to internalize and present antigen.

Within secondary lymphoid organs, APCs display intact, multivalent antigens on their surfaces for B cells to surveil (12–16). Using actin-based membrane protrusions, B cells rapidly probe APC surfaces with their BCRs (17). This process typically lasts just five minutes when cognate antigen is absent (12, 15). Upon encountering cognate antigen, a B cell establishes a longer-lived immune synapse, characterized first by actin-driven spreading across the APC surface to maximize antigen binding, followed by coordinated actions of the actin and microtubule networks to sweep antigens from the synapse periphery to its center (18–20). During this centripetal movement, actomyosin structures generate pulling forces on BCR-antigen bonds, rupturing low-affinity interactions while extracting high-affinity antigens for internalization (21). This process selectively promotes the survival of high-affinity B cells (22), ultimately driving the production of high-affinity antibodies (23).

B cells use mechanical forces not only to extract membrane-presented antigens but also to sense physical properties of APCs. B cells engage APCs through a chain of noncovalently linked proteins including the BCR, antigen, tethering molecules (e.g., antibodies and complement), and APC receptors (e.g., Fc and complement receptors) (24, 25). Forces propagated from BCRs through these molecular connections create a tug-of-war that makes antigen acquisition sensitive to the relative strength of each binding interface (26). For example, when B cells pull on antigens firmly anchored to stiff APC membranes, they achieve precise affinity discrimination but limited antigen extraction (27). Conversely, weakly tethered antigens or soft APC membranes enhance extraction efficiency at the expense of affinity discrimination (27). Thus, the precision of B cell antigen capture depends not only on BCR-antigen bond affinity and mechanical resistance, but also on the physical properties of APCs and their antigen-anchoring molecules.

APC membranes vary not only in stiffness but also fluidity, which is governed by membrane viscosity and interactions between membrane molecules and the cortical cytoskeleton (28). These biophysical characteristics determine antigen mobility on the APC surface, profoundly influencing early events in B cell activation. High antigen mobility enhances BCR microcluster growth, signaling, and transport (29, 30), whereas low antigen mobility promotes cell spreading (30) and traction force generation (31, 32). Yet, the impact of antigen mobility on B cell antigen internalization remains unexplored. It is plausible that highly mobile antigens are preferentially internalized through enhanced activation of signaling pathways that assemble force-generating actomyosin structures (33). Alternatively, frictional coupling of low-mobility antigens to centripetal actin flow via the BCR may load higher forces that are more capable of wrenching antigens from their tethers (34, 35). Moreover, mobility-dependent force variations in the synapse could influence both the spatiotemporal dynamics of antigen extraction and the subsequent trafficking of internalized antigens to processing compartments.

To explore these possibilities, we used DNA nanostructures to create multivalent antigens and quantify their force-mediated extraction from model membranes of different viscosity. We hypothesized that increasing membrane viscosity, thereby reducing antigen mobility, would amplify forces on BCR-antigen bonds and enhance the efficiency and precision of antigen uptake. Our results strongly support this model and reveal an unexpected temporal advantage: reduced antigen mobility accelerates these processes, enabling B cells to make activation decisions within physiologically relevant timescales. These findings suggest that APCs, by presenting immobilized, multivalent antigens, may actively shape antibody responses by promoting the rapid and selective activation of high-affinity B cells.

## Results

### Controlling antigen valency and mobility with DNA nanostructures and planar lipid bilayers

B cell activation outcomes are influenced by both the valency and mobility of antigens (30, 36, 37). To independently control these parameters, we combined antigen-coupled DNA nanostructures with glass-supported planar lipid bilayers. DNA nanostructures provide precise control over the number of linked antigens, unlike protein scaffolds that offer only statistical control over antigen multimerization (38). Additionally, a defined number of fluorophores can be added to each DNA structure for quantitative analysis of antigen densities (39).

Each DNA structure was covalently bound to three copies of NIP (4-hydroxy-3-iodo-5-nitrophenyl), a hapten antigen that binds the BCR of primary naïve B1-8 B cells with a 3D K_d_ value of 0.33 µM (21) (Fig. 1A). The N-hydroxysuccinimide-functionalized haptens were attached to the DNA via Uni-Link modifications, featuring amino groups separated from the phosphodiester backbone by a four-carbon chain (Fig. S1A). Due to the high conformational flexibility of these moieties, we computationally analyzed the structure to determine the mean inter-hapten distance, which ranged from <1 nm at equilibrium to 2.9 nm when fully extended (Fig. S1B). Given that the IgM-class BCR cannot bind bivalently to antigens spaced closer than 3.6 nm (40), each trivalent antigen-DNA construct likely induces crosslinking of 2-3 BCRs.

**Figure 1.**
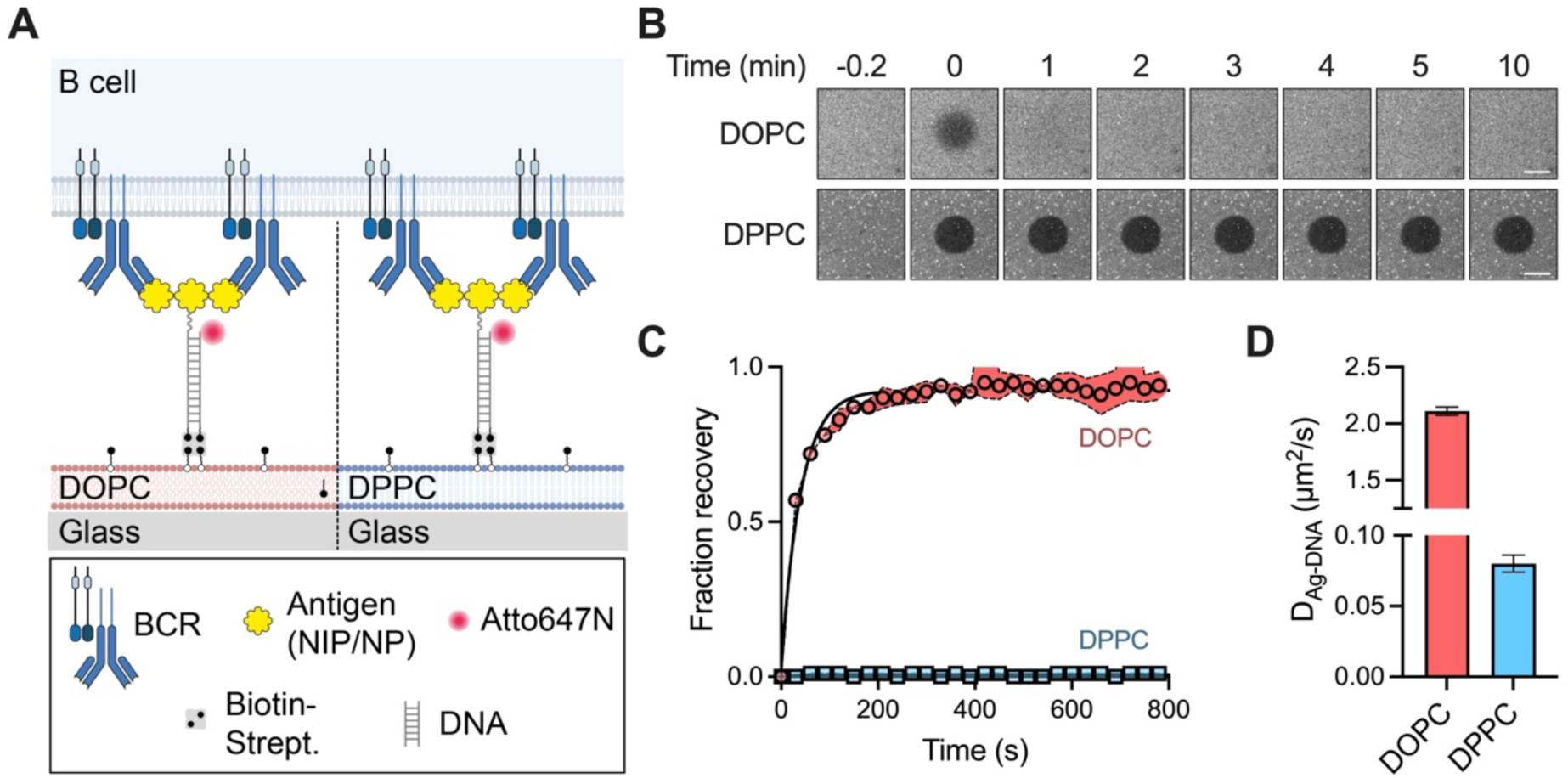
Changing antigen mobility by changing substrate viscosity. (A) Schematic showing BCR crosslinking by DNA-conjugated antigens tethered to glass-supported lipid bilayers. (B-D) Membrane fluidity characterization. (B) Time-lapse fluorescence images showing FRAP of labeled lipids. Scale bar: 20 µm. (C) FRAP recovery curves (mean ± SEM) from DOPC and DPPC bilayers, with a minimum of three regions analyzed per bilayer. (D) Diffusion coefficients of antigen-DNA sensors on each bilayer type, determined by single-particle tracking. Data from 673 (DOPC) and 1857 (DPPC) trajectories across three independent experiments. Bars represent mean ± SEM.

To manipulate antigen mobility, we constructed planar lipid bilayers (PLBs) using either DOPC or DPPC lipids. These lipids have identical head groups, maintaining a constant chemical composition for interfacing with cells, but different tail groups that alter lipid packing, resulting in surface viscosities of 8.4 × 10^−11^ Pa·s·m for DOPC and 3.0 × 10^−9^ Pa·s·m for DPPC (41). Fluorescence recovery after photobleaching (FRAP) measurements of PLBs doped with fluorescent lipids confirmed the contrasting viscosities of DOPC and DPPC bilayers (Fig. 1B). DOPC bilayers recovered to 95% with a half-time of 28 s, as expected for highly mobile lipids in a low-viscosity membrane, while DPPC bilayers recovered by just ∼2%, indicative of a high-viscosity membrane that limits lipid movement (Fig. 1C).

To quantify the impact of bilayer viscosity on antigen mobility, we attached antigen-DNA structures to the PLBs using biotin-streptavidin interactions and measured their diffusion by single-particle tracking (Fig. 1D and Movies S1 and S2). The mean diffusion constants were 2.1 ± 0.04 µm^2^/s on DOPC and 0.08 ± 0.006 µm^2^/s on DPPC, indicating that a 35-fold increase in PLB viscosity resulted in a 26-fold decrease in antigen-DNA mobility (Fig. 1D). This change agrees with a Stokes-Einstein-like expression (42) relating the diffusion constant, *D*, to the bilayer surface viscosity, *η_m_*, as

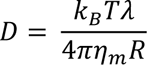

where *k_B_* is the Boltzmann constant, *T* is the absolute temperature, *λ* is the characteristic length indicative of membrane perturbation (on the order of a single lipid, 0.5 nm) (41), *R* is the radius of the diffusing particle (43), and *η_m_* = 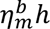, where 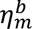 is the bulk viscosity of the bilayer and ℎ the bilayer thickness (5.9 nm for DOPC and 6.3 nm for DPPC) (41, 44). For our experiments, we assume + to be the radius of two lipids (1 nm), as each antigen-DNA construct is doubly biotinylated and tethered to biotinylated lipids in the bilayer through a tetravalent streptavidin molecule. Of note, the diffusion constant of antigen-DNA on the DPPC bilayer closely matches the mean diffusion constant of immune complexes on subcapsular sinus macrophage surfaces (0.011 µm^2^/s) (45). Thus, by altering bilayer viscosity, we can reliably influence antigen diffusion to mimic mobility characteristics that B cells encounter *in vivo*.

### High substrate viscosity limits antigen binding but enhances cell spreading and signaling

To investigate how bilayer viscosity—and consequently, antigen mobility— affects early B cell activation events, we stimulated B1-8 B cells using NIP_3_-DNA-coated PLBs. The NIP_3_-DNA density was adjusted to ∼3000 per µm^2^ on average (Table S1), corresponding to ∼9000 NIP per µm^2^. This density is comparable to antigen presentation on follicular dendritic cells, which express ∼300 complement receptor 2 molecules per µm^2^ that present multivalent antigens such as HIV virions (each expressing ∼10 trimeric envelope proteins) (46, 47). For these experiments, we used a DNA duplex with a shear force application geometry that cannot be ruptured by the forces B cells exert on BCR-antigen bonds (Fig. 1A; Table S2) (22, 48).

Using total internal reflection fluorescence (TIRF) microscopy, we observed that cells spread more extensively on DPPC than on DOPC bilayers (Fig. 2, A and B), indicating that B cells exert higher traction forces on surfaces with greater viscosity (41). Though increased spreading has previously been associated with increased antigen binding (18), we observed that B cells formed smaller BCR-antigen microclusters and accumulated less antigen overall on DPPC compared to DOPC, as indicated by the mean (Fig. 2C) and total antigen intensity (Fig. 2D), respectively. On DOPC, B cells clustered large amounts of antigen at the center of the cell-bilayer contact, as typically observed on these fluid surfaces. In contrast, on DPPC, antigens remained distributed throughout the cell-bilayer interface and were less clustered.

**Figure 2.**
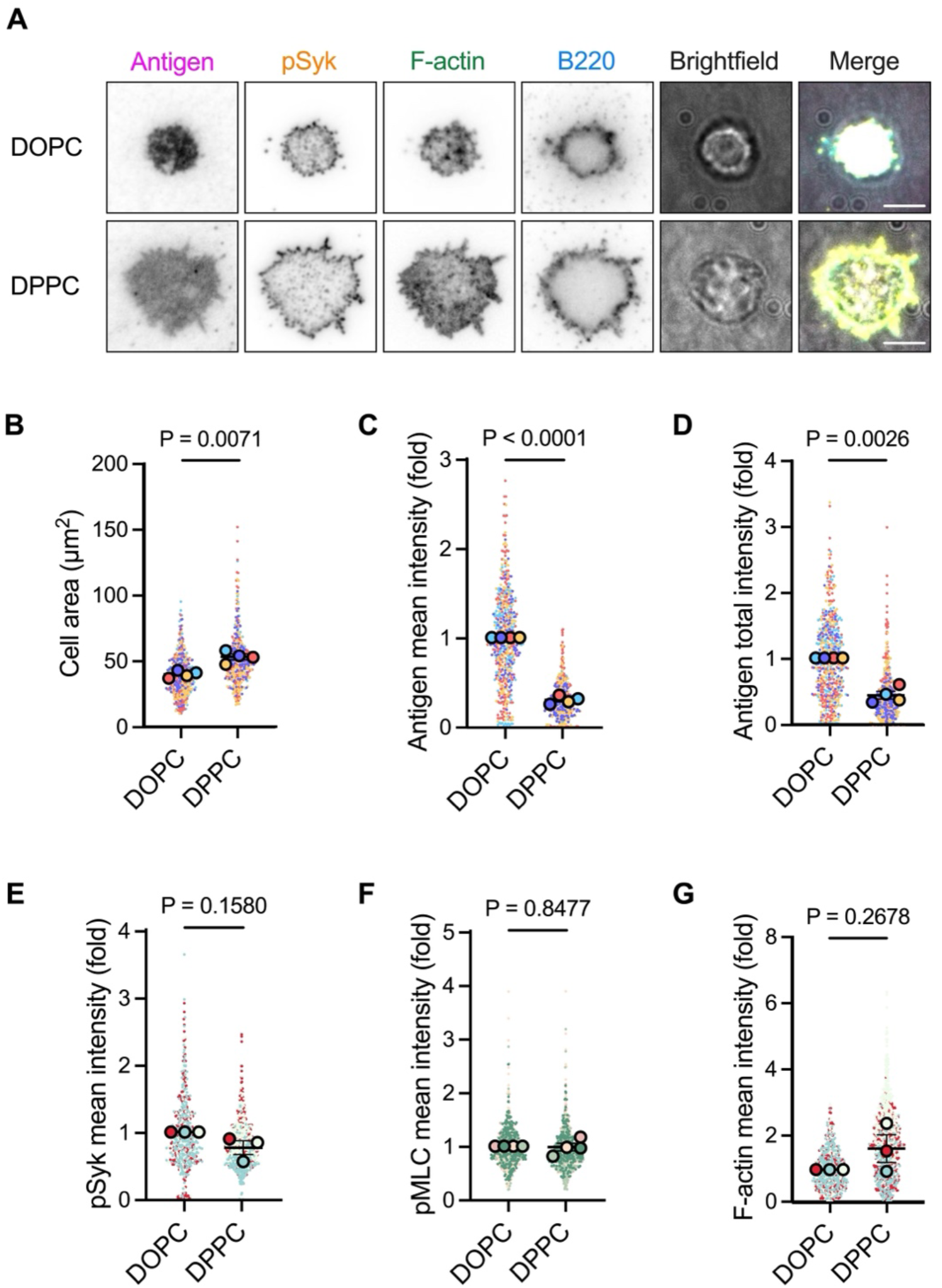
Lowering antigen mobility limits antigen binding but enhances B cell spreading and signalling. (A) TIRF microscopy images showing antigen, phosphoproteins, and filamentous (F)-actin distribution in B220+ B1-8 B cell synapses formed on DOPC (top) and DPPC (bottom) planar lipid bilayers after 15 minutes stimulation with NIP_3_-DNA. Scale bars: 5 µm. (B-G) Quantification of synaptic parameters: cell spreading area (B), mean (C) and total (D) antigen accumulation measured at 45 min, and levels of phospho-Syk (pSyk) (15 min) (E), phospho-myosin light chain (pMLC) (10 min) (F), and F-actin (G) (15 min). Individual cells (solid dots) and experiment means (outlined dots) are shown, color-coded by experiment. Data represent 3-4 independent experiments with cell numbers: B-D (DOPC n=533, DPPC n = 444, 4 experiments); E, G (DOPC n=521, DPPC n=888, 3 experiments); F (DOPC n=534, DPPC n=602, 4 experiments). Bars represent mean ± SEM. P values were determined by paired t-tests comparing experiment means.

Despite binding less antigen on DPPC, B cells phosphorylated Syk (Fig. 2E) and myosin light chain (Fig. 2F), and polymerized actin (Fig. 2G), as effectively as they did on DOPC. This suggests that BCR crosslinking by a trivalent antigen may be sufficient to induce robust signaling even in the absence of global BCR remodeling in the synapse (38), perhaps by bringing the local antigen concentration above a critical threshold needed for activation (49). Additionally, as the BCR is a mechanosensitive receptor (50), the higher traction forces facilitated by increased surface viscosity may amplify BCR signaling to compensate for reduced antigen binding.

### Calcium flux and transcription factor translocation occur effectively across both substrate viscosities

The recruitment and phosphorylation of signaling proteins close to the BCR lead to the mobilization of intracellular calcium, which encodes information through its amplitude and oscillations (51). This, in turn, activates transcription factors such as Nuclear Factor Kappa B (NF-κB) and Nuclear Factor of Activated T cells (NFAT) to regulate long-term cellular processes including survival, proliferation, and differentiation (51, 52).

We assessed intracellular calcium mobilization by loading B cells with the fluorogenic calcium-sensitive dye Cal-520 AM, seeding them onto NIP_3_-DNA-coated bilayers, and imaging them live at 37 °C. Imaging and quantification revealed that B cells responded to stimulation on DOPC with intracellular calcium fluxes slightly higher than those of B cells stimulated on DPPC (Fig. 3A), though the difference was not statistically significant (Fig. 3B). The calcium fluxes occurred with similar kinetics, as measured by the time delay between the cell first touching the bilayer and reaching maximum Cal-520 intensity (Fig. 3C).

**Figure 3.**
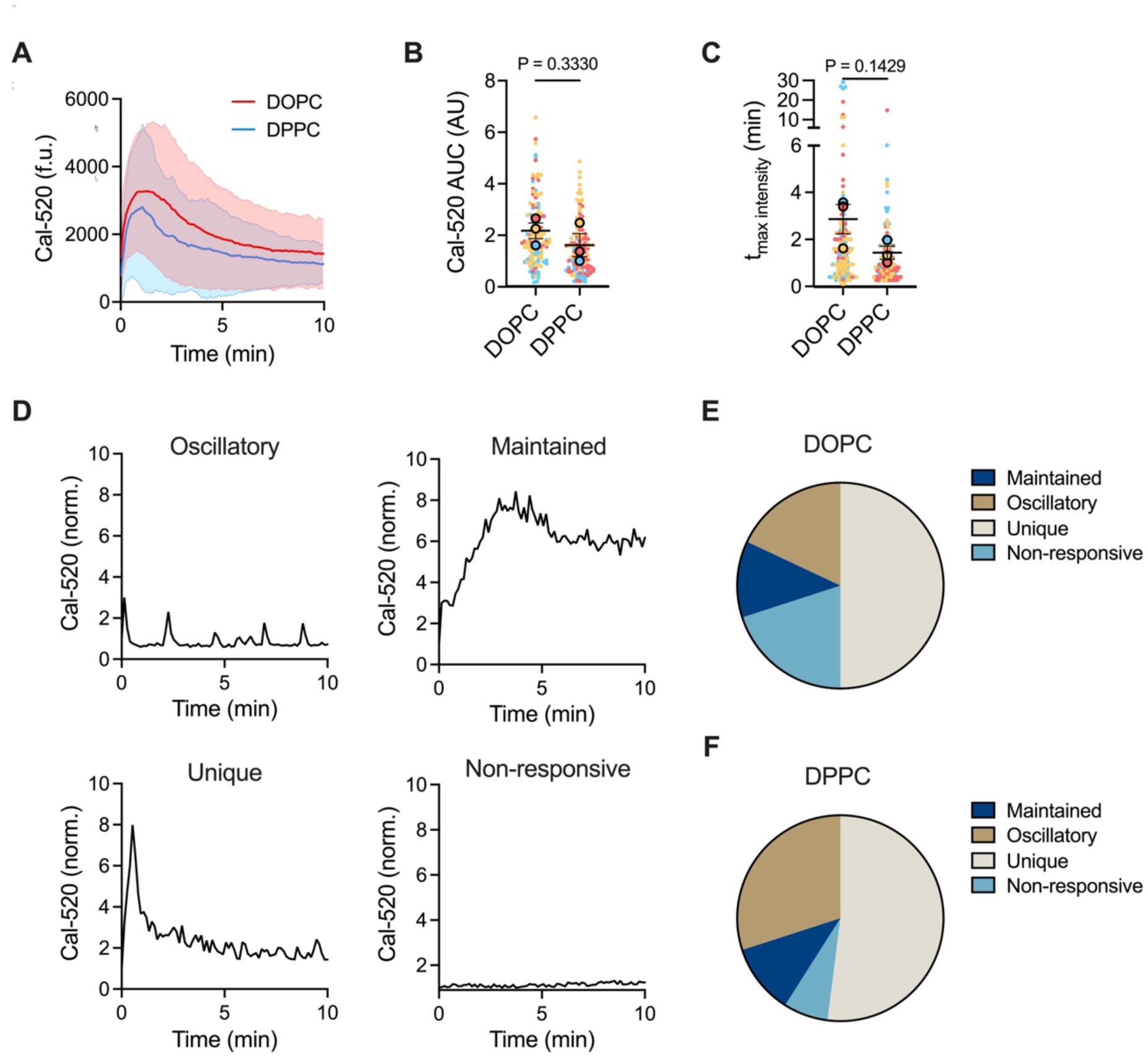
Antigen mobility does not impact B cell calcium signaling. (A-F) Analysis of calcium signaling in B1-8 naïve B cells responding to NIP_3_-DNA sensors. (A) Intracellular calcium levels over time on DOPC versus DPPC bilayers. Mean ± SD shown. (B) Total calcium response quantified as area under the curve (AUC, arbitrary units). (C) Time delay between initial BCR-antigen binding and peak calcium signal. (D) Representative calcium traces showing four distinct response patterns: oscillatory, maintained, unique, and non-responsive. (E,F) Distribution of calcium response patterns on DOPC (E) and DPPC (F) bilayers. Data from 135 (DOPC) and 127 (DPPC) cells across three independent experiments. Individual cells (solid dots) and experiment means (outlined dots) are shown, color-coded by experiment. Bars in (B, C) represent mean ± SEM. P values were determined by paired t-tests comparing experiment means.

We also analyzed calcium oscillation patterns, categorizing the flux for each cell into one of four categories: oscillatory, maintained, unique, and non-responsive (Fig. 3D). Cells were more likely to flux calcium on DPPC (93%) than on DOPC (80%) but otherwise showed broadly similar patterns, with the greatest fraction of cells having a single (unique) peak (Fig. 3, E and F). Correspondingly, B cells stimulated by both surfaces translocated NF-κB component p65 (Fig. 4, A and B) and NFAT (Fig. 4C) to the nucleus to the same extent. When defining activation as nuclear-to-cytoplasmic ratio >1, approximately 66% of cells activated NF-κB and 82% of cells activated NFAT. These results indicate that both high- and low-mobility antigens elicit robust signal propagation through calcium release to NF-κB and NFAT, resulting in similar BCR-triggered signaling phenotypes.

**Figure 4.**
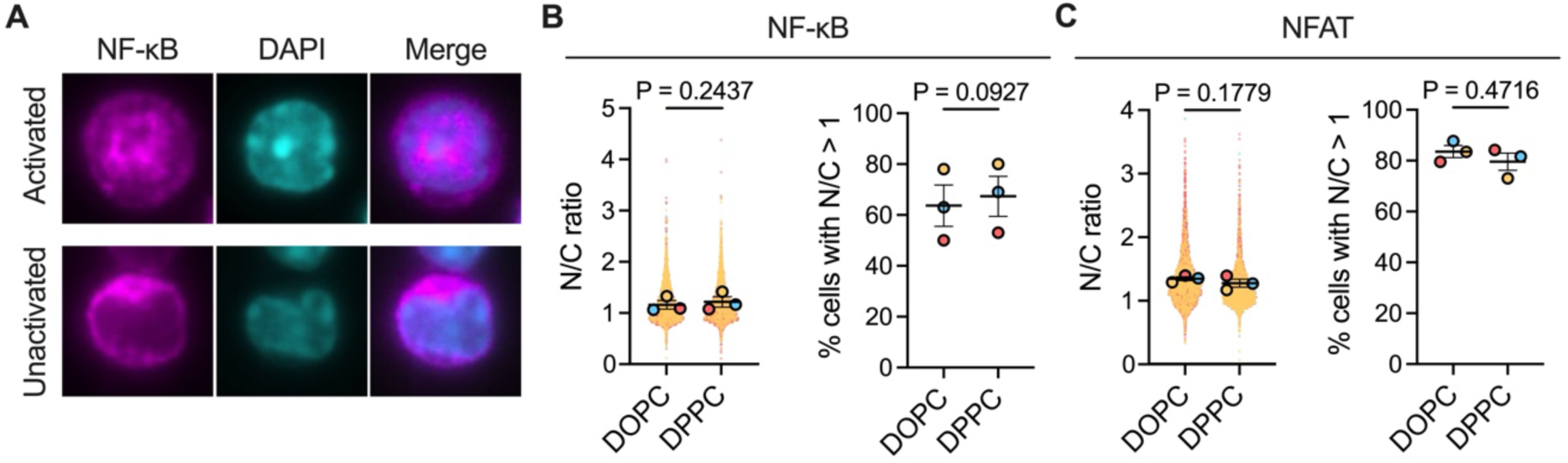
The activation of NF-κB and NFAT transcription factors is unaffected by antigen mobility. (A) Activation is measured on a cell-by-cell basis by calculating the nuclear-to-cytoplasmic ratio of transcription factor intensity. Top: active state with nuclear translocation; bottom: inactive state with cytoplasmic retention. Scale bars: 5 µm. (B, C) Quantification of NF-κB (B) and NFAT (C) activation, showing nuclear-to-cytoplasmic intensity ratios (I_nuc_/I_cyt_; left) and percentage of activated cells (I_nuc_/I_cyt_ > 1; right). Data are from three independent experiments: NF-κB (DOPC n=6231; DPPC n=4221) and NFAT (DOPC n=4296; DPPC n=3498). Individual cells (solid dots) and experiment means (outlined dots) are shown, color-coded by experiment. Bars represent mean ± SEM. P values were determined by paired t-tests comparing experiment means.

### B cells extract antigens more efficiently from high-viscosity substrates

The previous results suggested that substrate viscosity has little impact on BCR signaling in response to trivalent antigens. However, the reproductive fitness of a B cell during an immune response is primarily determined by the amount of antigen that B cells extract and internalize in the immune synapse. Therefore, we next focused on this aspect of B cell activation.

To quantify antigen extraction, we used a sensor containing a DNA duplex that can be unzipped to release NIP_3_ from the surface under an applied force of about 10 pN (Fig. S2) (27). The sensor features an Atto647N fluorophore and Iowa Black RQ quencher pair at the base of the duplex. Upon duplex rupture, the Atto647N becomes fluorescent and is transported along with the NIP_3_ antigen into endosomal compartments. Since each NIP_3_ antigen is transported with one Atto647N fluorophore, the Atto647N fluorescence intensity serves as a proxy for the total amount of antigen captured by the cell. To quantify the total amount of antigen bound in the synapse, the bottom DNA strand—remaining on the substrate after duplex rupture—is labelled with an Atto550 fluorophore, which is not affected by the quencher.

B cells accumulated less antigen in synapses formed on DPPC than on DOPC after 45 minutes of stimulation (Fig. 5A). This was assessed by quantifying the mean (Fig. 5B) and total (Fig. 5C) Atto550 signal intensity in the synapse and is consistent with earlier results using non-internalizing NIP_3_ antigens (Fig. 2, C and D). Despite binding substantially less antigen on DPPC bilayers, B cells internalized comparable amounts relative to cells on DOPC, as evidenced by the total antigen extracted (Fig. 5D), the number of extracted antigen clusters (Fig. 5E), and the antigen quantity per cluster (Fig. 5F) (all evaluated in the Atto647N channel). Consequently, B cells internalize a larger proportion of available antigen molecules in synapses formed on DPPC bilayers (Fig. 5G), indicating that high-viscosity substrates enhance the efficiency of antigen capture by B cells.

**Figure 5.**
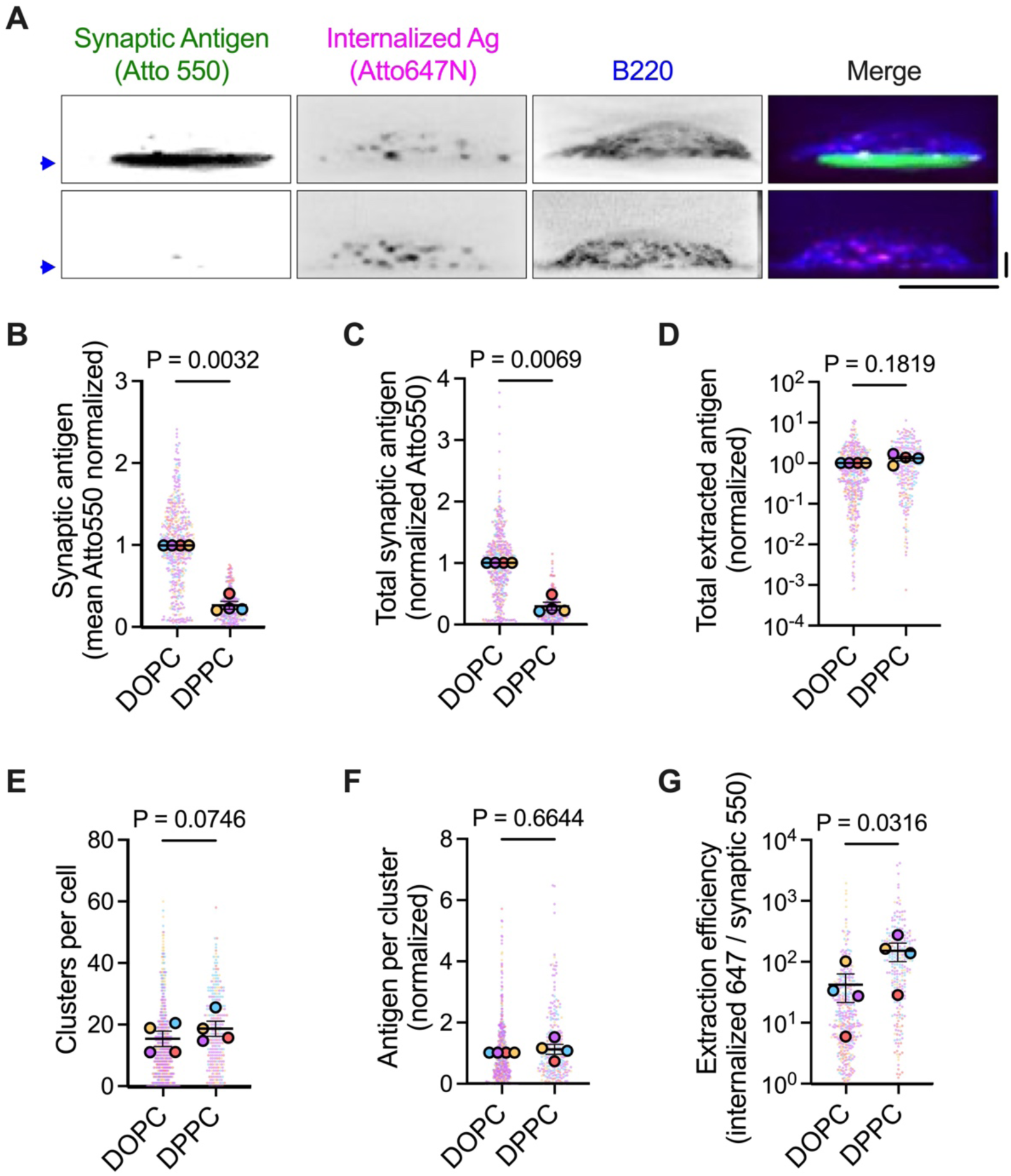
B cell antigen extraction is more efficient when antigens have low mobility. (A) Side-view reconstructions of B220+ B1-8 B cells showing membrane-bound (Atto550) and internalized (Atto647N) NIP_3_ antigen on DOPC and DPPC bilayers. Blue arrows indicate the bilayer positions. Scale bars: 5 µm. (B, C) Synaptic antigen accumulation after 45 min, quantified as mean (B) and total (C) intensity. (D-F) Characterization of extracted antigen clusters: total antigen extracted (D), clusters per cell (E) and antigen intensity per cluster (F). (G) Extraction efficiency was calculated as the ratio of internalized (Atto647N) to bound (Atto550) antigen intensity per cell. Data are from 472 (DOPC) and 264 (DPPC) cells across four independent experiments. Individual cells (solid dots) and experiment means (outlined dots) are shown, color-coded by experiment. Bars represent mean ± SEM. P values were determined by paired t-tests comparing experiment means.

### Substrate viscosity influences the spatiotemporal dynamics of antigen extraction in the synapse

While measuring antigen extraction after 45 minutes reveals B cells’ capacity to acquire antigens from presenting membranes, it does not capture the dynamics of this process. Given the transient nature of B cell contacts with APCs (12, 15), we sought to determine whether differences in antigen mobility affected the kinetics of antigen capture. This was possible because mechanical rupture of the DNA duplex leads to Atto647N unquenching and the appearance of distinct fluorescent spots that can be detected and tracked by live-cell microscopy (Fig. S3; Movie S3). By quantifying the time between a cell first contacting the bilayer and internalizing its first antigen cluster, we found that B cells internalized antigen significantly faster from DPPC (mean: 74 s) compared to DOPC (mean: 421 s) (Fig. 6A). Further, 64% of B cells internalized at least one antigen cluster from DPPC within 5 minutes, compared to only 26% from DOPC (Fig. 6B). These results indicate that lowering antigen mobility enables B cells to capture antigens on the fast timescales of B cell-APC interactions *in vivo*.

**Figure 6.**
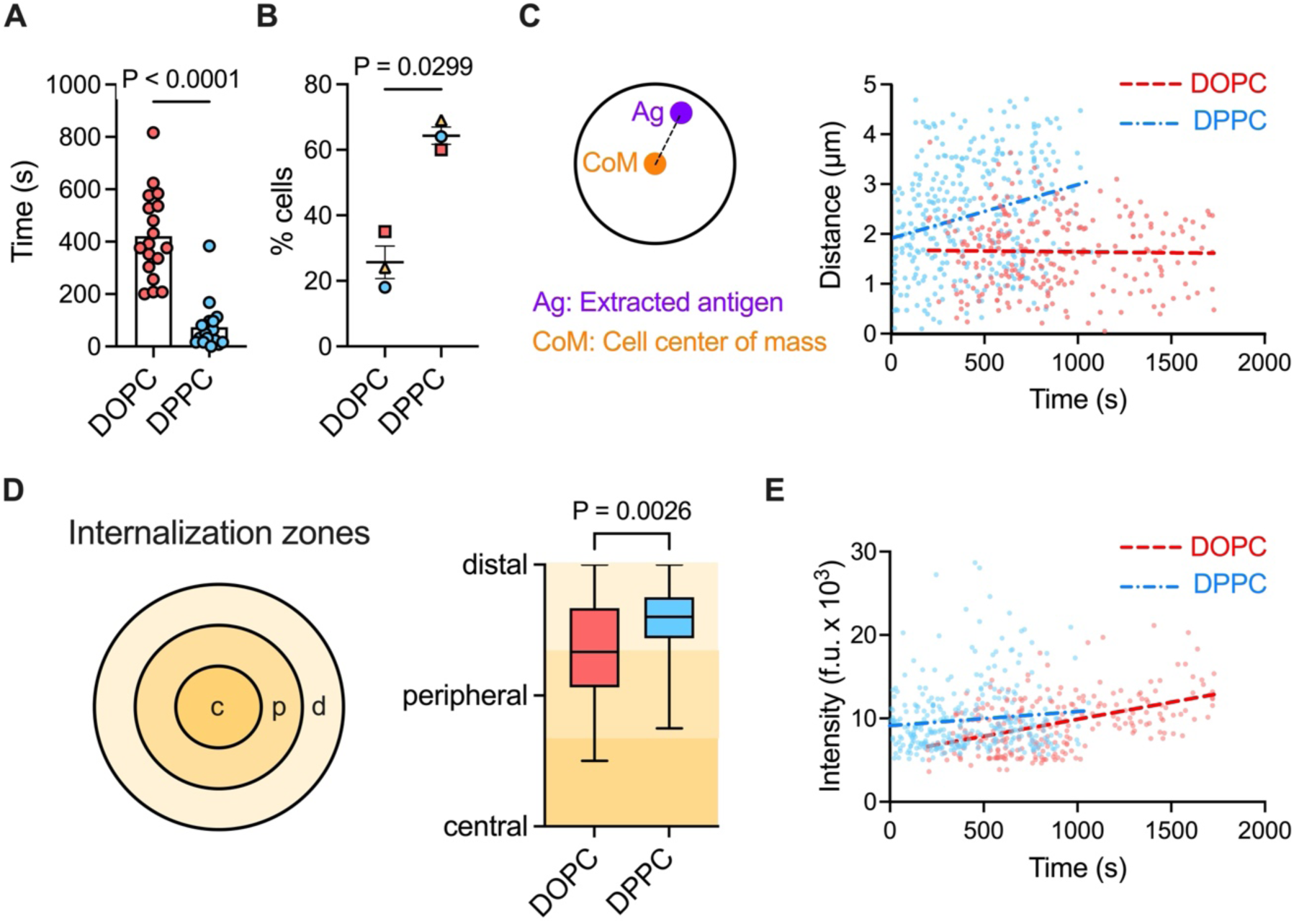
B cells extract antigens with different spatiotemporal dynamics from DOPC and DPPC bilayers. (A) Time to first antigen cluster extraction after initial bilayer contact. Individual cells are shown. (B) Percentage of cells extracting at least one antigen cluster within 5 min of contacting the bilayer. The points show experiment means and bars represent mean ± SEM. (C) Relationship between extraction time and distance from the cell center. Pearson correlation analysis shows a positive correlation on DPPC (r = 0.3, p < 0.0001) but not on DOPC (r = −0.02, p = 0.77). (D) Distribution of extraction events across central, peripheral, and distal synaptic regions. Boxes indicate the quartile range (first quartile to third quartile), the horizontal line indicates the median value, and the whiskers extend to the minimum and maximum points. (E) Cluster size (Atto647N intensity) versus extraction time. Pearson correlation analysis shows a positive correlation on both substrates: DOPC (r = 0.47, p < 0.0001), DPPC (r = 0.11, p = 0.0457). Data represent 245 (DOPC) and 324 (DPPC) antigen clusters extracted by 18 cells per condition, pooled from three independent experiments. Each dot in C and E represents a single extracted cluster.

A current hypothesis suggests that the centralization of antigens within the synapse is necessary for their extraction and internalization by B cells (53). To test this, we analyzed the spatial coordinates of extracted antigen clusters from single-particle trajectories, measuring the distance of each cluster from the cell center using an anti-IgM Fab surface label to define the cell boundary (Fig. 6C, left). A plot of distance versus time revealed distinct spatiotemporal dynamics of antigen extraction from DOPC and DPPC bilayers (Fig. 6C, right). On DPPC, B cells initially internalized antigens at a mean distance of 1.9 µm from the cell center, increasing to 3 µm by 17 minutes. In contrast, on DOPC, internalization consistently occurred at a mean distance of 1.7 µm from the cell center over 30 minutes. This suggests that antigen mobility influences the location of extraction within the synapse.

Given that B cells spread differently on DOPC and DPPC bilayers, we further analyzed the data by quantifying the fraction of antigen clusters extracted from the central, peripheral, and distal regions of the synapse (Fig. 6D, left). These regions were calculated for each frame to account for changing cell area over time. The results showed that antigen extraction is skewed toward the cell edge, with >80% of clusters on DOPC and >90% of clusters on DPPC extracted from either the peripheral or distal regions (Fig. 6D, right). Only 15% on DOPC and 7% on DPPC were extracted from the synapse center. Thus, centralization is not required for B cells to capture antigens from presenting membranes.

B cells cluster antigens to different extents in synapses formed on DOPC and DPPC bilayers (Fig. 2A, C, and D). We therefore investigated whether the size of extracted antigen clusters changes over time to reflect variations in antigen accumulation in the synapse. We quantified the intensity of each extracted antigen cluster in its initial detection frame and plotted intensities against extraction time (Fig. 6E). A positive correlation was found on both DOPC (r = 0.47, p < 0.0001) and DPPC (r = 0.11, p = 0.0457), indicating that antigen clustering in the synapse promotes the extraction of larger antigen clusters. This effect was more pronounced on DOPC, reflecting the greater extent of antigen clustering on this highly mobile substrate.

### Internalized antigens are transported to MHCII+ compartments consistently across varying substrate viscosities

It is crucial for the antibody response that antigens are delivered to MHCII+ compartments for processing and presentation. The observation that B cells capture antigens from DOPC and DPPC bilayers with distinct spatial and temporal kinetics led us to question whether the antigens may be trafficked differently within the cell. To investigate this, we allowed B cells to interact with NIP_3_-DNA-coated bilayers for 20 minutes and then fixed and stained them intracellularly with an anti-MHCII antibody. Z-stack images and quantification revealed that internalized antigen clusters were trafficked to MHCII+ compartments (Fig. 7A), as assessed both by the intensity of anti-MHCII staining per internalized antigen cluster (Fig. 7B) and the percentage of internalized antigen clusters that colocalized with MHCII (Fig. 7C). These findings suggest that despite differences in the efficiency, dynamics, and location of antigen internalization, B cells direct internalized antigens to the correct intracellular compartments for processing and presentation.

**Figure 7.**
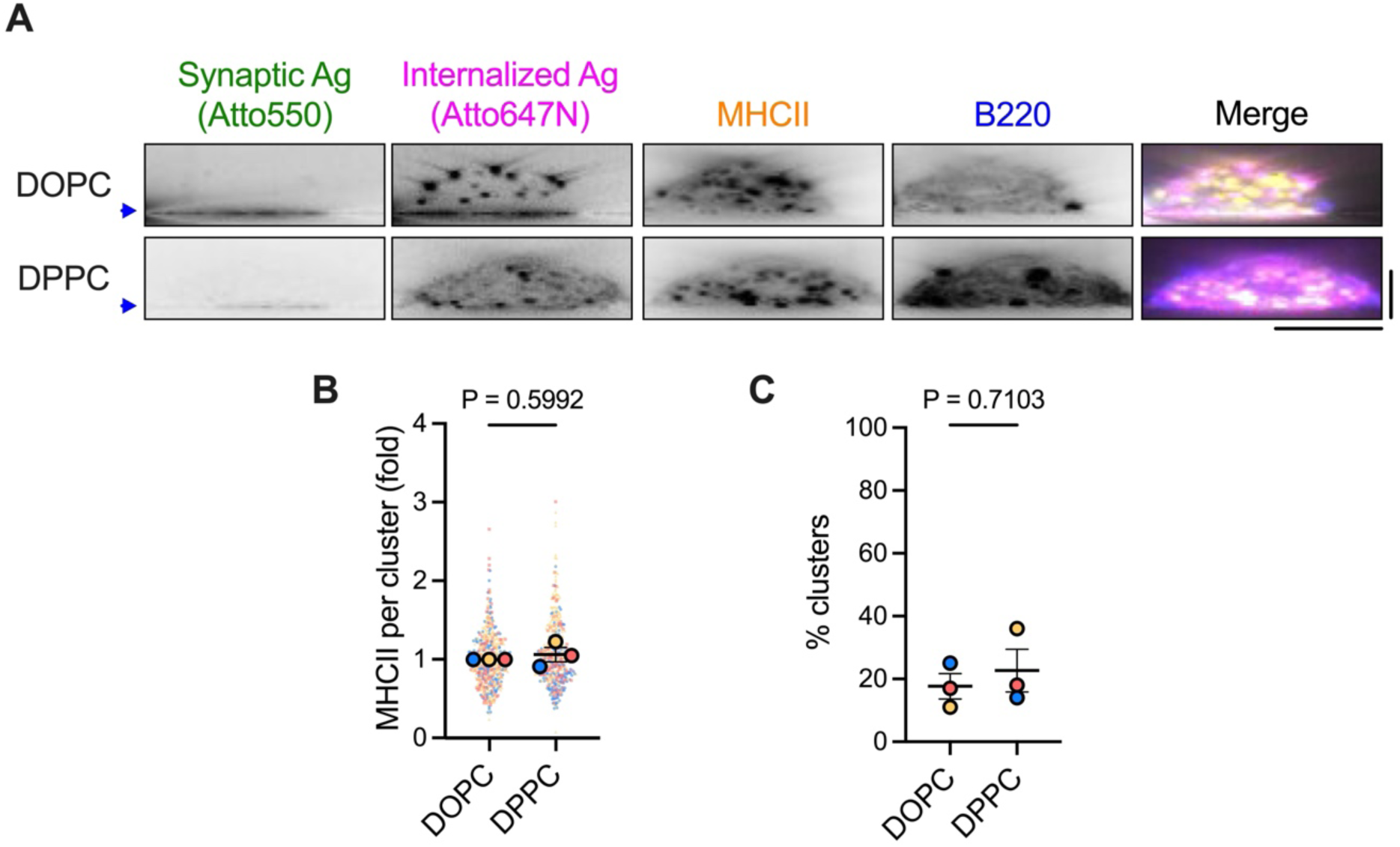
B cells traffic antigens extracted from DOPC and DPPC membranes to MHCII+ compartments at the same rate. (A) Side-view reconstructions showing B220+ cells with internalized NIP_3_ antigen clusters colocalizing with MHCII. The blue arrows indicate bilayer positions. Scale bars: 5 µm (horizontal), 10 µm (vertical). (B, C) Quantification of MHCII+ trafficking: the mean MHCII intensity per internalized cluster (B) and the percentage of MHCII+ clusters (C). Data are from 538 (DOPC) and 475 (DPPC) cells across three independent experiments. Individual cells (solid dots) and experiment means (outlined dots) are shown, color-coded by experiment. Bars represent mean ± SEM. P values were determined by paired t-tests comparing experiment means.

### High substrate viscosity enhances B cell discrimination of antigen affinities

We previously demonstrated that substrate stiffness affects the forces applied to BCR-antigen bonds, influencing B cells’ ability to discriminate antigen affinities (27). To determine if antigen mobility has a similar effect, we compared the ability of B cells to distinguish NIP_3_ from NP_3_ (4-hydroxy-3-nitrophenyl), a hapten that binds the B1-8 Fab with ∼10-fold lower affinity (21). After 45 minutes, B cells had discriminated NIP_3_ and NP_3_ on DPPC, as shown by cell spread area (Fig. 8A), antigen binding (Fig. 8B), and Syk phosphorylation (Fig. 8C). On DOPC, discrimination was more modest, primarily reflected as a difference in antigen binding (Fig. 8B). The strongest effects were observed in antigen extraction, where B cells internalized significantly more NIP_3_ than NP_3_ from both substrates, measured by the number of internalized clusters per cell (Fig. 8D) and the percentage of cells acquiring at least one antigen cluster (Fig. 8E). However, the NIP_3_ to NP_3_ extraction ratio was significantly higher on DPPC (Fig. 8F), indicating that B cells achieve more stringent affinity discrimination when antigens have low mobility.

**Figure 8.**
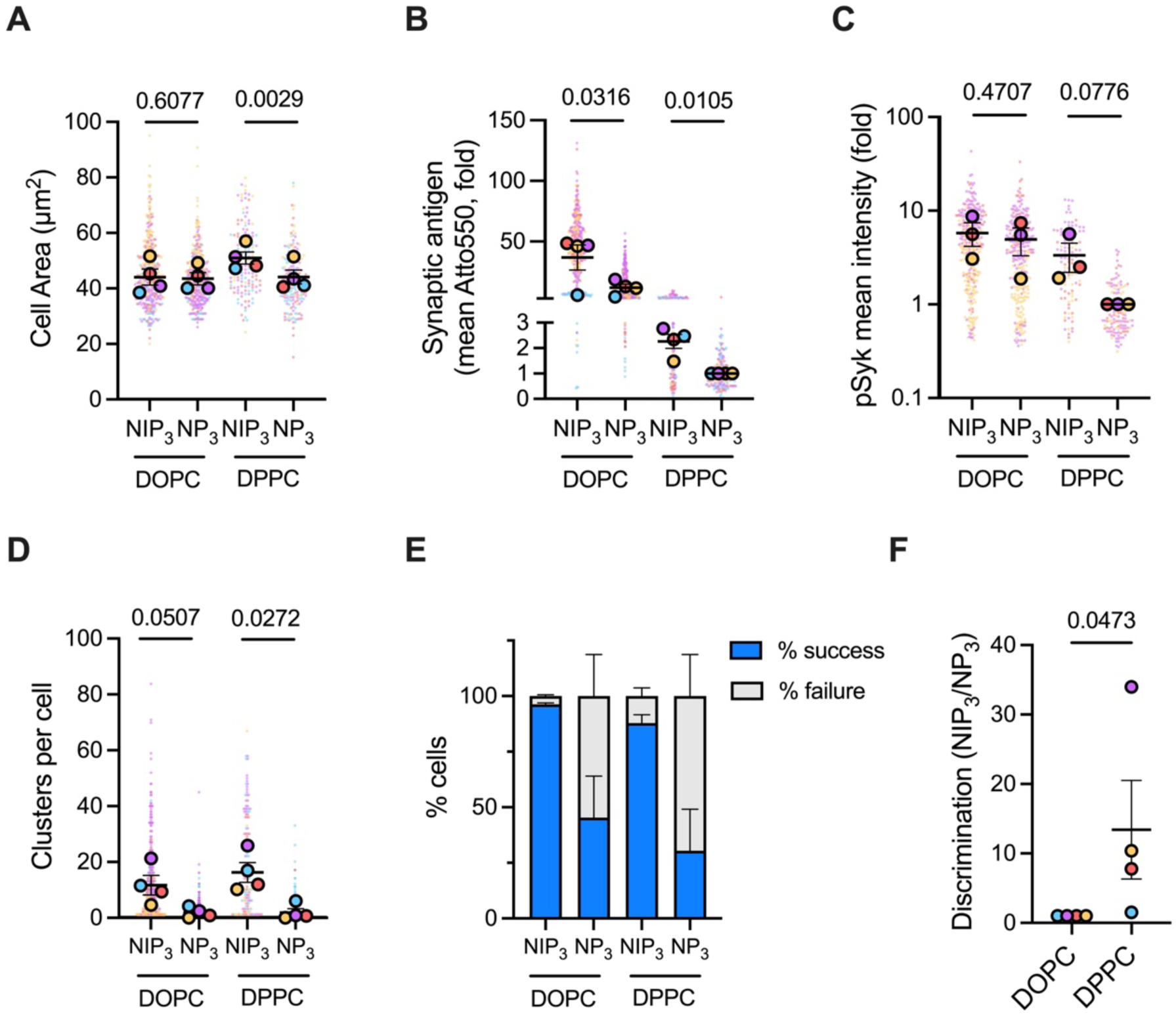
B cell antigen extraction is more stringent from high-viscosity substrates. (A-C) Quantification of cell spreading area (A), antigen clustering (B), and phospho-Syk signaling (C) in synapses formed on DOPC versus DPPC bilayers. (D, E) Effect of substrate viscosity on the number of internalized clusters (D) and the percentage of cells internalizing at least one cluster (E). (F) The ratio of NIP_3_ to NP_3_ antigen internalized from DOPC and DPPC. Data are from four independent experiments: DOPC (NIP_3_ n=410, NP_3_ n=368) and DPPC (NIP_3_ n=145, NP_3_ n=185). pSyk (C) was measured in three of the four experiments. Individual cells (solid dots) and experiment means (outlined dots) are shown, color-coded by experiment. Bars represent mean ± SEM. P values were determined by paired t-tests comparing experiment means.

## Discussion

B cells use mechanical energy to physically extract antigens from the surfaces of other cells (21). This process serves two purposes: internalizing antigens for processing and presentation (24), and discriminating antigens based on the force-dependent lifetimes of BCR-antigen bonds (54). Our study reveals that antigen mobility regulates the magnitude, location, and timing of forces in the immune synapse thereby regulating the speed and accuracy of B cell antigen capture and discrimination.

The actin cytoskeleton plays a central role in B cell activation, orchestrating immune synapse formation through rapid self-remodeling and consequent cell surface restructuring in response to BCR signaling. Actin polymerization at the leading cell edge drives cell spreading (18), while membrane tension together with myosin IIa contractility generate a retrograde actin flow that facilitates BCR-antigen clustering, signaling, and centripetal transport (55). Our study confirms previous observations that antigen mobility influences these actin-dependent behaviors, with highly mobile antigens enhancing clustering and transport, and low-mobility antigens promoting cell spreading (30). However, unlike earlier studies using monovalent antigens (29, 30), we observed no enhancement in proximal BCR signaling with increased antigen mobility. This extended to downstream processes, including calcium flux and the activation of NFAT and NF-kB transcription factors. These findings align with our recent work characterizing B cell activation by subcapsular sinus macrophages (56), suggesting that antigen mobility does not alter B cell fitness. The discrepancy with earlier work likely stems from our use of antigen trimers, which can crosslink multiple BCRs to initiate robust signaling (38). Antigen trimers feature in various pathogens, including viral spike proteins (e.g., HIV-1, SARS-CoV-2, influenza hemagglutinin) (57–59) and bacterial proteins (e.g., OmpF, TAAs) (60, 61), potentially representing an optimal antigenic unit for eliciting robust B cell activation across diverse physical environments.

Actin remodeling and myosin II contractility not only reshape the immune synapse but also exert mechanical forces on BCR-antigen bonds. Using antigen extraction as a proxy for force generation, we demonstrate that synaptic forces exhibit distinct spatiotemporal patterns depending on antigen mobility. Antigens presented on fluid-phase DOPC membranes are most likely extracted in the peripheral synapse region, where myosin II motors contract actin filaments to generate traction forces (20, 31, 32). In contrast, antigens displayed on gel-phase DPPC membranes are mainly extracted in the distal synapse region, where forces arise from rapid retrograde actin flow (20, 62). While we observe some antigen extraction in the central area of the synapse, consistent with previous studies (32, 33), this accounts for only a minor fraction of total antigen uptake. This suggests that flowing actin filaments, rather than dynamic actin puncta, predominate during antigen capture in the B cell synapse.

The coupling between actin flow and cellular adhesion complexes is a well-described mechanism for force transmission from a cell to its substrate (35, 63). These forces are, in turn, influenced by the substrate’s mechanical properties (41, 64, 65). When B cells pull on antigens embedded in a fluid membrane, much of the energy dissipates due to antigen and substrate remodeling that limits the shear forces that B cells can apply through retrograde actin flow. As a result, B cells primarily rely on tensile forces exerted by myosin II motors in the synapse periphery. Conversely, low-mobility antigens largely retain their structure within gel-phase membranes, directing more mechanical energy to BCR-antigen bonds and allowing B cells to leverage the high actin flow velocities in the distal region of the synapse for antigen extraction. Restricted growth of antigen clusters on gel-phase substrates also increases the mechanical load on individual BCR-antigen bonds (22, 66, 67). Higher forces not only increase the probability of antigen extraction through enhanced rupture of soft DNA tethers (26), but also shorten and narrow bond lifetime distributions (68), leading to improved antigen affinity discrimination and faster antigen capture. These results suggest that antigens of different mobility engage distinct cytoskeletal structures within the immune synapse, altering the fidelity of B cell selection.

That only ∼20% of internalized antigens are observed within MHCII+ compartments in our experiments highlights a need to further investigate how antigen mobility and synaptic patterning influence downstream B cell outcomes. It is possible that our fixed time-course imaging measurements capture only a fraction of the total antigens in these compartments as they traffic through complex and time-dependent routes required for processing and MHCII presentation (69). An alternative explanation is that B cells internalize antigens through different pathways depending on substrate mobility. B cells use at least two endocytic routes—clathrin-mediated endocytosis (21) and fast endophilin-mediated endocytosis (70)—for internalizing membrane-presented antigens. The proportion of antigens directed toward each pathway could be influenced by antigen cluster size (71) or mechanical tension (72), both of which we demonstrate are affected by antigen mobility. This could have important implications for B cell fate, as these distinct internalization routes not only provide metabolic support for B cell growth and proliferation (70), but also shape the peptide repertoire presented on MHCII (69). Further investigation into how antigen mobility affects antigen trafficking and presentation could provide valuable insights into B cell activation and immune function.

Our hybrid live cell-supported membrane platform has revealed that antigen mobility directly influences the speed and accuracy of naïve B cell activation. This finding is physiologically relevant, as APCs use their actin cytoskeleton to restrict antigen diffusion and cluster growth (56), potentially enhancing the activation of high-affinity B cells over lower affinity cells by maximizing forces on BCR-antigen bonds. However, the situation is complex. B cell antigen discrimination is also sensitive to APC membrane stiffness (27), which varies across cell types and is modulated by inflammatory signals (73). APCs can dynamically adapt their physical properties in response to changing mechanics and biochemical composition of the extracellular matrix (56), which is remodeled during the adaptive immune response (74, 75). Furthermore, B cells themselves undergo significant changes during immune responses, differentiating into germinal center and memory B cells with class-switched BCRs and distinct cytoskeletal architectures that alter mechanotransduction (22, 76, 77). This suggests that factors beyond BCR affinity are likely to influence the antibody repertoire, potentially helping to explain recent findings that germinal centers support the maturation of B cells with a breadth of affinities to yield diverse antibody responses (78, 79).

## Acknowledgements

We thank the KCL Biological Services Unit, and Kheya Sengupta and Laurent Limozin for helpful discussions. This work was supported by a BBSRC research grant (BB/S007814/1) and a BBSRC sLoLa (BB/V003518/1) to KMS.

## Author Contributions

H.C.W.M. and K.M.S. designed research; H.C.W.M. and M.I. performed research; H.C.W.M. and A.T.B. contributed new analytic tools; H.C.W.M. and K.M.S. analyzed data; H.C.W.M. and K.M.S. wrote the paper.

## SI Materials and Methods

### Mice

B1-8^flox/flox^ Igκ^Ctm1Cgn/tm1Cgn^ mice on a C57BL/6 background (B1-8 mice) were used as a source of primary naïve B cells for all experiments. All mice were between 14 and 22 weeks old and both males and females were used. Mice were bred and treated in accordance with guidelines set by the UK Home Office and the King’s College London Ethical Review Panel.

### Primary B cell isolation and culture

For each experiment, primary splenocytes were obtained by passing a mouse spleen through a 70-µm cell strainer and lysing red blood cells with RBC lysis buffer (Invitrogen). Cells were pelleted by centrifugation for 7 minutes at 300x*g* and 4 °C, resuspended in 450 µl ice-cold MACS buffer (PBS, pH 7.3, 0.5% BSA, 1 mM EDTA), and incubated with 50 µl CD43 (Ly-48) MicroBeads, mouse (Miltenyi Biotec) for 20 minutes on ice. The cells were resuspended in 10 ml ice-cold MACS buffer, pelleted, resuspended in 500 µl ice-code MACS buffer, and isolated by negative selection using an LD column and MidiMACS Separator (Miltenyi Biotec). Cells were pelleted and resuspended to a density of 5 x 10^6^ cell/ml in full RPMI (RPMI 1640 medium supplemented with 10% FBS, 1% MEM non-essential amino acids, 2 mM L-glutamine, 50 µM 2-mercaptoethanol, 100 U/ml penicillin, and 100 µg/ml streptomycin, all from Gibco), and cultured at 37 °C with 5% CO_2_ for 1-2 h before using in experiments.

### Preparation of glass coverslips

Glass coverslips (24 x 50 mm, No. 1.5; Fisher) were etched in piranha solution (2:1 H_2_SO_4_:30% H_2_O_2_) for 15 minutes, washed 10 times with ultrapure water (Sartorius Arium), and rinsed three times with 100% ethanol. Sample chambers were assembled by placing a strip of 10 µl CultureWell gaskets (Grace Bio-Labs) onto the coverslip, which was then glued onto a one-well Nunc Lab-Tek chamber (Thermo Fisher Scientific) using Sylgard fast-cure silicone (Silmid).

### Antigen-conjugated DNA tension sensors

The sensors were assembled as previously described (1). Briefly, HPLC-purified single-stranded DNA oligos (Tables S2 and S3) were purchased from Integrated DNA technologies and resuspended to 100 µM in duplex buffer (30 mM HEPES, pH 7.5, 100 mM potassium acetate; IDT). To assemble the sensors, oligos were mixed in stoichiometric amounts, supplemented with 2 mM MgCl_2_, heated to 95 °C for 3 minutes in a thermocycler, and cooled on the bench top to room temperature for 30 minutes. The annealed sensor was then exchanged into degassed 0.1 M sodium carbonate buffer, pH 8.3, using a 7-kDa molecular weight cut-off (MWCO) desalting column (Zeba; Thermo Fisher Scientific) and incubated with a 20-fold molar excess of either NIP-Osu (4-hydroxy-3-iodo-5-nitrophenylacetic active ester; Biosearch Technologies) or NP-Osu (4-hydroxy-3-nitrophenyl hapten active ester; Biosearch Technologies) per Uni-link modifier for 1 hour at room temperature to label the amine functional groups. The solution was then passed twice through desalting columns to remove unreacted hapten. A conjugation ratio of 3:1 was confirmed by UV-Visible spectroscopy (NanoDrop One^C^; Thermo Fisher Scientific) using the Atto550 (rupturable sensor) or Atto647N (non-rupturable sensor) absorbance as a proxy for DNA concentration. The sensors were stored at –20 °C in single-use aliquots.

### Estimating inter-hapten distance

Computational analysis of the trivalent antigen followed a two-step approach combining Avogadro and CREST calculations. The initial fully extended structure was generated in Avogadro (2) to estimate the maximum inter-hapten distance. To determine minimum inter-hapten distance, we first optimized the structure using xTB-6.6.1, using the semiempirical tight binding-based quantum chemistry method GFN-xTB with implicit water solvation (3). We then used CREST-2.12 to identify the lowest energy conformer through computational sampling (4). The simulation was run for 0.5 ps using the GFN2-xTB method with implicit water solvation, maintaining all other parameters at default values. Molecular structures from these analyses (Fig. S1) were visualized using CYLview20 (C. Y. Legault, Université de Sherbrooke, http://www.cylview.org), with inter-hapten distances measured in Avogadro.

### Planar lipid bilayers

Small unilamellar vesicles (SUVs) were prepared by mixing either 97% 1,2-dioleyol-*sn*-glycero-3-phosphocholine (DOPC) or 97% 1,2-dipalmitoyl-*sn*-glycero-3-phosphocholine (DPPC), 2% 1,2-dioleoyl-sn-glycero-3-[(N-(5-amino-1-carboxypentyl)iminodiacetic acid) succinyl] (nickel salt) (DGS-NTA(Ni)), and 1% 1,2-dioleoyl-sn-glycero-3-phosphoethanolamine-N-(cap biotinyl) (biotin-DOPE; Avanti Polar Lipids, Inc.) in chloroform at a final lipid concentration of 4 mg/ml (5). The solvent was dried with a gentle stream of argon and then under vacuum for at least 3 h. The lipid film was resuspended to 5 mM in degassed PBS at room temperature (DOPC) or degassed SUV buffer (DPPC; 10 mM Tris, pH 7.5, 150 mM NaCl) pre-warmed to 70 °C by vortexing and then bath sonicating until the suspension cleared (about 1 h) to produce SUVs. The SUVs were centrifuged for 10 min at 16,000x*g* and 4 °C (DOPC) or 40°C (DPPC) to remove large particulates. The SUVs were stored under argon at 4 °C and used to prepare bilayers for a maximum of 1 month.

Planar lipid bilayers were prepared by diluting DOPC SUVs to 0.2 mg/ml in PBS, and DPPC SUVs to 0.2 mg/ml in fusion buffer (10 mM Tris, pH 7.5, 300 mM NaCl, 10 mM MgCl_2_) pre-warmed to 70 °C, and adding 10 µl to a CultureWell gasket attached to a piranha-etched coverslip. SUVs were incubated on the coverslip for 45 min in a sealed humidity chamber at either room temperature (DOPC) or 70 °C (DPPC) to allow vesicles to fuse. The bilayers were washed with PBS (DOPC), or with fusion buffer followed by PBS (DPPC) to remove unfused vesicles. The bilayers were then incubated sequentially with 100 µg/ml BSA for 1 h, 20 min with 0.5 µg/ml streptavidin, and 10 min with antigen-conjugated DNA sensors, which were titrated to the densities reported in Table S1. Sensor densities were matched to within +/- 30% across all bilayers on the day of each experiment. To assess bilayer mobility, FRAP measurements were performed on bilayers doped with 0.005 mol % Liss Rhod PE lipids using an A1R+ confocal microscope (Nikon). A circular region of radius ∼17 µm was photobleached for 2 s using a 561-nm laser modulated by a galvanometer scanner. Fluorescence recovery was monitored at 30 s intervals using a gallium arsenide phosphide cathode (GaAsP) detector.

### Quantification of DNA sensor density on bilayers

The surface density of fluorescent tension sensors was calibrated following the procedure developed by Galush et al (6). DOPC bilayers were doped with either Liss Rhod PE or Cy5.5 PE lipids at concentrations ranging from 0 to 0.06 mol % or 0 to 0.5 mol %, respectively, and imaged with the same conditions used for cell-based measurements to generate a calibration curve to map fluorescence intensity to number of lipid fluorophores. To use the calibration curve to determine the number of antigen molecules on the bilayer, the sensor fluorescence intensity was compared to the lipid fluorescence intensity to obtain the F factor, defined as: F = I_bulk(sensor)_/I_bulk(lipid)_, where I_bulk(sensor)_ and I_bulk(lipid)_ are the intensities of DNA tension sensor or lipid in solution at the same concentration. The intensity values were measured in imaging buffer, 2 µm above the glass coverslip.

### Fluorescence microscope

TIRF and z-stack images were acquired using a Nikon TiE TIRF microscope equipped with a 100x, 1.49-NA oil-immersion objective (Nikon), a motorized stage with an integrated piezo Z-drive (MS-2000; Applied Scientific Instrumentation), and active Z-drift correction (Perfect Focus System; Nikon). The microscope was controlled by a high-speed TTL, I/O, DAC controller (Triggerscope 4; Cairn Research) integrated into MicroManager software (7). Illumination was supplied by a MultiLine LaserBank (Cairn Research) fitted with 405-, 488-, 561-, and 640-nm diode lasers (Coherent OBIS). The beams were aligned into a single-mode fibre and coupled to an iLas2 Targeted Laser Illuminator (Gataca Systems), which produces a 360° spinning beam with an adjustable illumination angle. Laser beams were passed through a laser quadband (405/488/561/640nm) filter set for TIRF applications (TRF89901-v2-ET; Chroma) before illuminating the sample. Emitted photons were filtered by appropriate single-band emission filters (Chroma) using a filter wheel (OptoSpin; Cairn Research) and then captured onto a back-illuminated sCMOS camera (Prime 95B sCMOS; Teledyne Photometrics). For live-cell imaging, a relative humidity of 95% and a constant temperature of 37 °C was maintained using a cage incubator fitted with an active humidity controller (Okolab).

### Single-particle tracking analysis of antigen-DNA diffusion

Bilayers were incubated with a low density of antigen-DNA sensors to enable detection of single particles. The sensors were imaged using 561 nm excitation, with a power density of 18 W/cm^2^ at the sample, to visualise the Atto550 fluorophore labelling the DNA. Images were acquired with 10 ms exposure and no delay between frames at 37 °C. Sensors were detected and tracked using the ImageJ plugin TrackMate (8). A Difference of Gaussians filter was used to detect spots of ∼0.5 µm diameter using a quality threshold of 2 and sub-pixel localization. Simple LAP tracker was used for tracking, with a 1-µm linking maximum distance, 1.5-µm gap-closing maximum distance, and 2-frame or 20-ms gap-closing maximum gap. Trajectories with 9 or more steps were carried forward for mean-squared displacement (MSD) analysis.

The MSD for each trajectory was calculated using Track Processor MSD in the Track Manager plugin of Icy (9). For a Brownian particle diffusing in two dimensions,

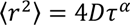

where ⟨*r*^2^⟩ is the displacement (µm^2^) of the particle during time interval *τ* (s), *D* is the diffusion constant (µm^2^/s), and *α* is the anomalous diffusion coefficient that defines Brownian (*α* = 1), sub-diffusive (*α* < 1), and super-diffusive (*α* > 1) motion. The diffusion constant was calculated using the first 26 time lags for DOPC and the first 11 time lags for DPPC, where the plots of ⟨*r*^2^⟩ versus *τ* were linear.

### Fixed-cell imaging

B cells were washed and resuspended in warm Hank’s balanced salt solution, calcium, magnesium, no phenol red (HBSS; Gibco) supplemented with 0.1% BSA (HBSS 0.1% BSA). Cells were allowed to recover at 37 °C for 5 min and then added to HBSS 0.1% BSA in pre-warmed imaging chambers with the antigen-coated bilayers and incubated for the indicated timepoints at 37 °C to allow B cells to form immune synapses and extract antigen for internalization. Cells were fixed in 2.6% paraformaldehyde (PFA) for 10 min, blocked with 5% (v/v) normal mouse serum (Jackson ImmunoReserach) for 30 min, and stained with 1 µg/ml anti-mouse/human B220 for 30 min at room temperature or overnight at 4 °C. Stained cells were washed, permeabilised, and then incubated with primary antibodies for intracellular staining (Table S4).

For staining with anti-NF-κB (p65) and anti-NFAT, cells were permeabilised with the FoxP3 fixation/permeabilization kit (BioLegend). For all other intracellular stains, cells were permeabilised with HBSS supplemented with 1% BSA, 5% normal mouse serum, and 0.3% Triton X-100. After washing with permeabilization buffer, cells were incubated with secondary antibodies and phalloidin to stain filamentous actin. Cells were PFA fixed and washed again before imaging either in TIRF or by acquiring z-stacks with a 0.5-µm step size.

To measure antigen affinity discrimination, B cells were stimulated on bilayers presenting either NIP_3_-DNA or NP_3_-DNA complexes. There were four conditions for each experiment: DOPC NIP_3_-DNA, DOPC NP_3_-DNA, DPPC NIP_3_-DNA, and DPPC NP_3_-DNA. B cells were stimulated in parallel across all substrates and imaged with the same imaging conditions.

### TIRF image processing

Images were cropped to remove poorly illuminated regions, background subtracted, and flatfield-corrected before proceeding to image analysis. Semi-automated ImageJ macros segmented cells based on B220 surface marker staining where possible and obtained quantitative information from signalling channels. Where B220 staining was not available, phalloidin staining was used to segment the cells. All segmentations were assessed, and poorly segmented cells were removed prior to analysis.

### Z-stack image processing

Prior to image processing and analysis, images were cropped, background-subtracted, and flatfield-corrected. Analysis of multi-color z-stack images was performed with a user-guided pipeline in MATLAB (MathWorks) tethered to ImageJ via the MIJ plugin (http://bigwww.epfl.ch/sage/soft/mij/) as previously described (10). Briefly, cells were detected first in 2D and then in 3D using both the B220 and brightfield channels. 3D cell masks were then stored as image stacks for subsequent analysis, excluding any cells touching the edge of the image. The synapse was identified as the sharpest image plane in the Atto550 channel. Antigen extraction was analysed for each cell by bandpass filtering each plane and identifying Atto647N-positive clusters above the synapse using a user-defined global threshold. Antigens were considered internalised if they were at least 1.5 µm above the synapse plane. Local background due to fluorescence scattered from the substrate was subtracted from each identified antigen cluster. Data were inspected and manually gated using a custom-written visualization tool in MATLAB to remove any non-cellular debris, incorrectly segmented cells, and cells poorly stained with anti-B220.

The mean and total Atto550 fluorescence intensity of the pixels in the synapse plane were used as a measure of synaptic antigen binding. The total extracted antigen was calculated as the sum of Atto647N pixel intensities in all extracted antigen clusters. The efficiency of antigen extraction was calculated as the total extracted antigen intensity (Atto647N) divided by the total synaptic antigen intensity (Atto550). The mean Atto647N intensity of extracted clusters was used as a measure of antigen per cluster. Masks of the extracted antigen clusters, synapse, and total cell were used to quantify fluorescence in other channels. Cell spread area was calculated from the area of the cell mask at the synapse plane. Cells were considered to have failed to extract antigen if no antigen clusters were detected in the cell.

### Transcription factor translocation

B cells were stimulated with activating bilayers for 1 h at 37°C, fixed and stained with anti-B220 to label the cell surface, permeabilised and stained with either anti-NF-kB (p65) or anti-NFAT and DAPI, and imaged at a single plane approximately 2 µm above the bilayer surface where the DAPI staining was sharpest. Images were analysed in CellProfiler (11). For each cell, the nucleus and cytosol were identified from the DAPI and B220 channels and masked as different regions, and the transcription factor mean intensity quantified in each region. The nuclear translocation for each cell was calculated as the ratio of nuclear-to-cytosolic mean intensity.

### Live-cell calcium imaging and analysis

B cells were loaded with 5 µM Cal-520 AM (AAT Bioquest) at 37 °C in full RPMI for 1.5-2 h at a density of 5 x 10^6^ cells/ml. After 1.5-2 h, cells were washed into fresh RPMI and stained at the surface with Alexa Fluor 405 anti-IgM Fab (Jackson ImmunoResearch). For Fab staining, cells were stained for 30 min on ice at a density of 6 x 10^6^ cells/ml, with mixing by pipetting every 10 min (12). Cells were pelleted and resuspended in warm HBSS 0.1% BSA and allowed to recover for 5 min at 37 °C immediately before imaging. Cells were imaged on sensor-coated bilayers for at least 10 min by time-lapse TIRF microscopy, with images acquired every 8 s.

For analysis, cells were segmented using the anti-IgM Fab stain using K-means thresholding in a custom Icy pipeline to obtain the cell area and background-subtracted intracellular calcium intensity over time. Calcium traces were normalised to the first frame that the cell is in contact with the bilayer and were categorized based on the following rules. If the calcium intensity did not reach at least double the intensity value in frame 1, it was considered “non-responsive”. Otherwise, a line was drawn at the full width at half maximum (FWHM) for the first Ca^2+^ peak. If the calcium trace crossed the line twice, resulting in a single peak, it was categorized as “unique”; if the trace crossed the line four or more times, resulting in at least two peaks, it was categorized as “oscillatory”; if the trace crossed the line once and remained above it for the duration of the measurement, it was categorized as “sustained”.

### Live-cell imaging and analysis of antigen extraction

B cells were seeded onto sensor-coated bilayers and imaged sequentially in the 561-nm and 640-nm channels to visualise synaptic and extracted antigen, respectively. Cells were imaged for at least 10 minutes with images acquired every 8 s. The Atto647N fluorophore remains quenched in the synapse by Iowa Black RQ, and is unquenched only upon DNA tether rupture, indicating antigen extraction. Extracted antigen clusters were identified in widefield as spots having an intensity at least 1.5x the background Atto647N intensity within the cell.

For each experiment, 6 representative cells that internalized antigen from DOPC and DPPC substrates and had sharp antigen spots were selected for detailed analysis. A maximum projection of the anti-IgM Fab signal was used to segment the cell using default thresholding in ImageJ. The cell center of mass was then obtained using the Analyze Particles function. TrackMate (8) was used to detect and track the position and intensity extracted antigen clusters in the Atto647N channel. The xy coordinates of the first point in each trajectory were taken to be the position of antigen extraction and were used to calculate the distance from the cell center of mass. The time point of antigen extraction was taken to be the time between when the cell first contacted the bilayer and the time when the extracted antigen cluster appeared.

To bin extracted antigen clusters into the central, peripheral, or distal regions, at each time point in a live-cell video each cell was divided into three concentric circles of equal width centred around the cell centre of mass, which was identified based upon the anti-IgM surface stain. Extracted clusters were assigned to a bin based upon the xy coordinates of the frame in which they first appear.

### Figures and statistics

Graphs were plotted using SuperPlots formatting (13), where data for individual cells (small data points) and the mean values per experiment (large, outlined data points) were plotted together. Graphing and statistical analysis were performed using GraphPad Prism (versions 9 and 10). Where possible, exact P values are shown on the plots. P > 0.05 is considered non-significant (ns) and P ≤ 0.05 is considered significant (P ≤ 0.05 = *, P ≤ 0.01 = **, P ≤ 0.001 = ***, P ≤ 0.0001 = ****). A description of statistical tests and number of independent experiments is provided in the figure legends.

**Fig. S1.**
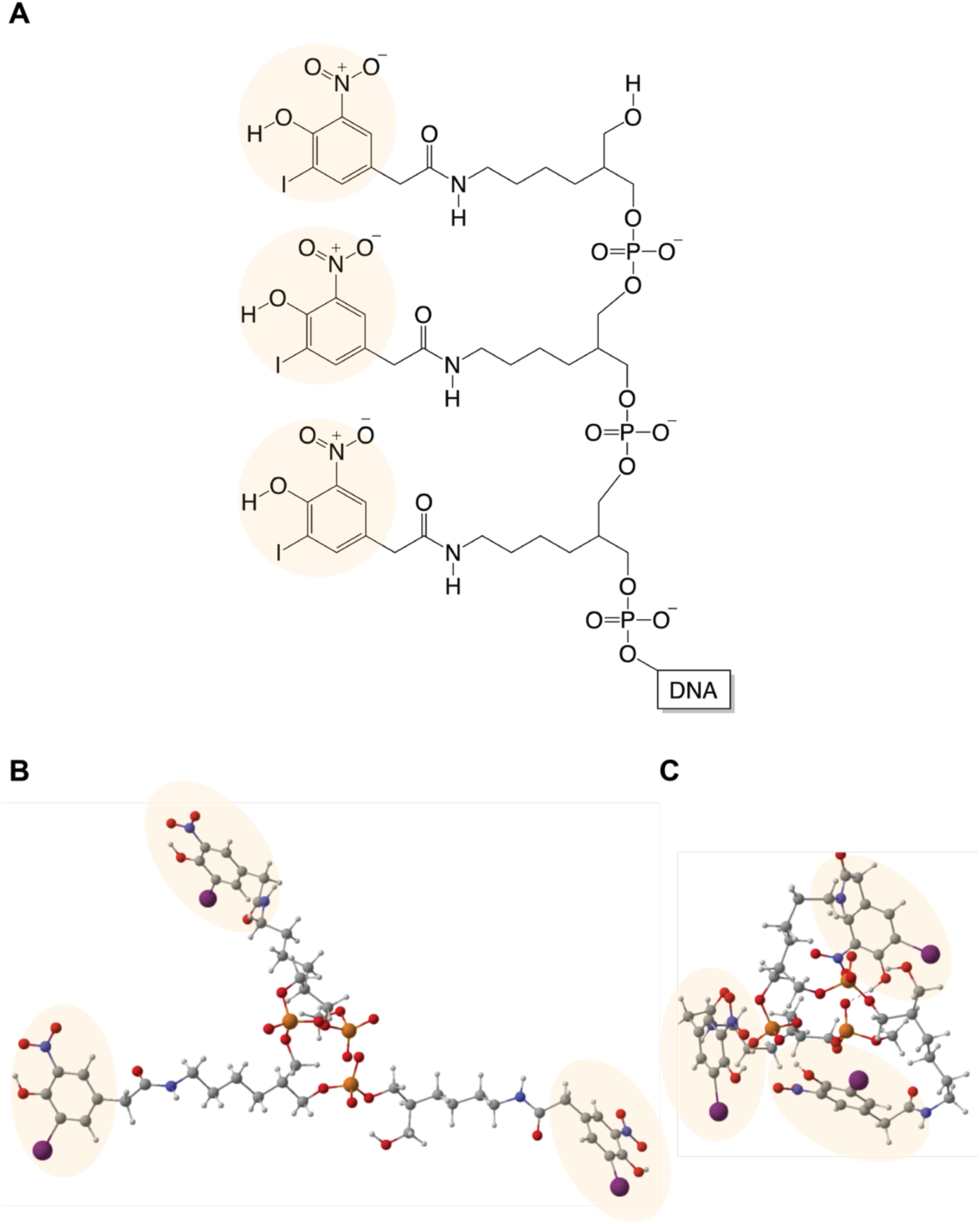
(A) Molecular structure of NIP_3_ antigen. (B, C) To compute the structures, the DNA molecule indicated in (A) was replaced with a methyl group. (B) The fully extended structure used to estimate the maximum inter-hapten distance. (C) The CREST-optimized structure used to estimate the equilibrium (minimum) inter-hapten distance. Atom color-coding: Gray-carbon; white-hydrogen; blue-nitrogen; red-oxygen; orange-phosphorus; purple-iodine. Hapten moieties are highlighted with yellow circles.

**Fig. S2.**
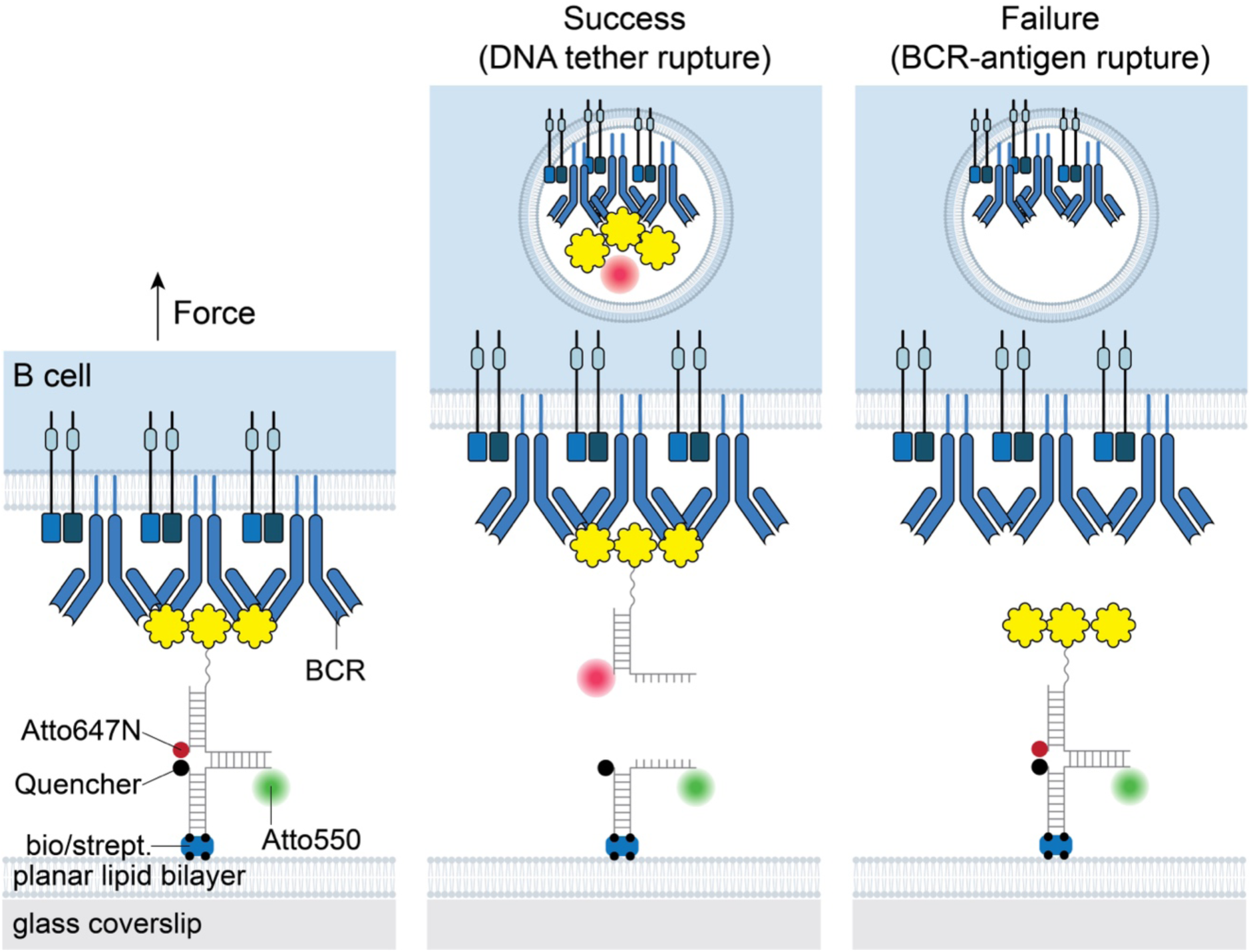
Schematic of the DNA tension sensor used to quantify force-mediated antigen extraction.

**Fig. S3.**
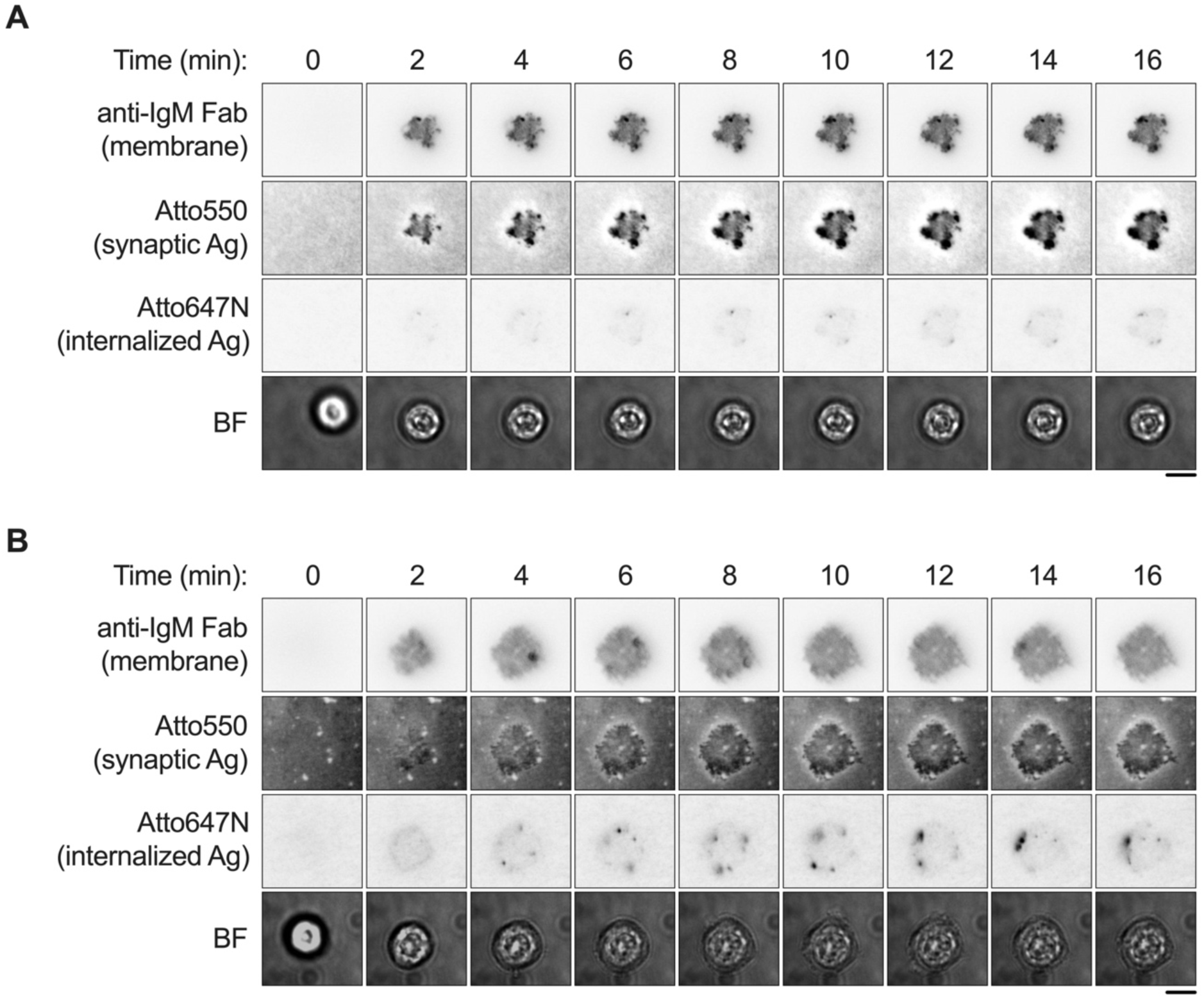
Still images of B cell antigen extraction from DOPC (A) and DPPC (B). Extracted clusters appear as dark spots in the Atto647N channel. Scale bars: 5 µm.

**Table S1.**
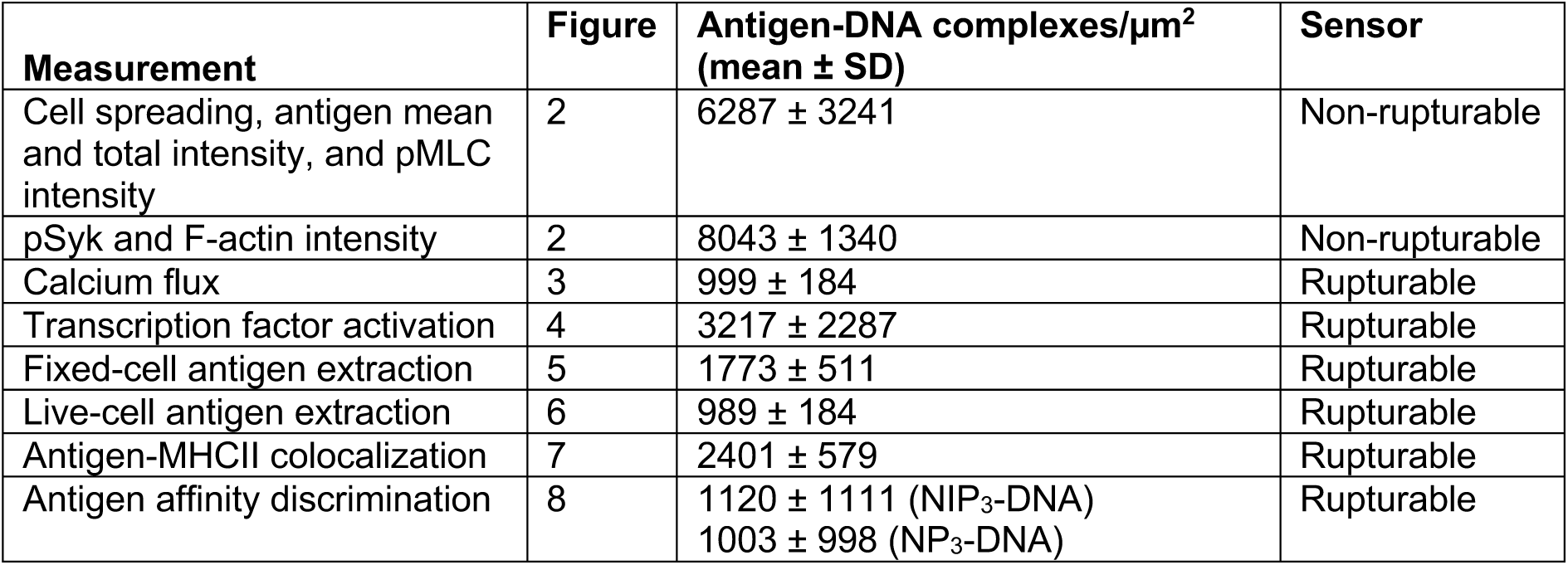
Antigen densities and sensors used for each experiment. The average surface density of antigen-DNA complexes across all experiments was 2870 ± 1506 (mean ± SD)

**Table S2.**
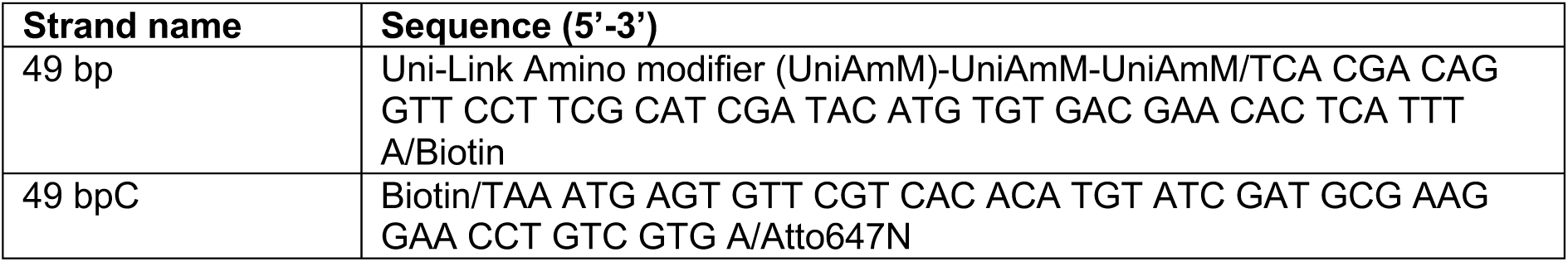
Oligonucleotide sequences used to make the non-rupturable DNA sensor.

**Table S3.**
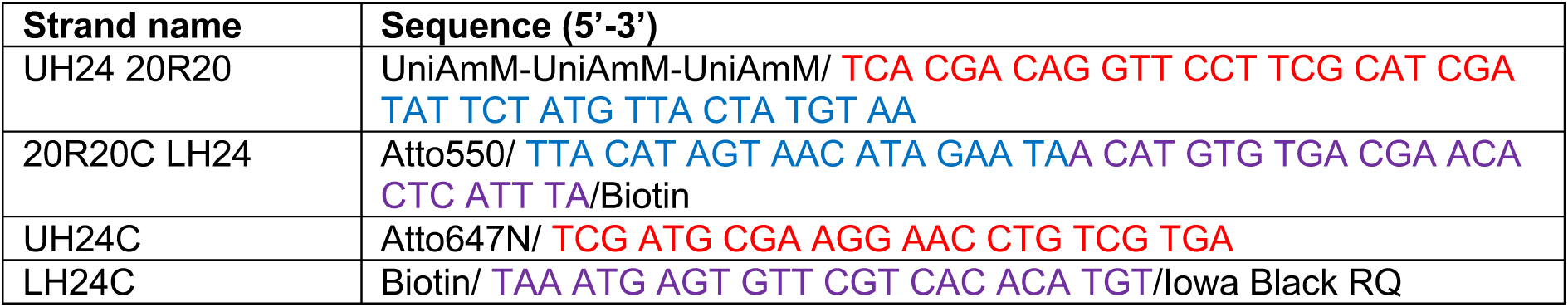
Oligonucleotide sequences used to make the rupturable DNA sensor. Color coding: red—24-bp upper handle; purple: 24-bp lower handle; blue: 20-bp tether.

**Table S4.** Antibodies and probes used for fluorescence staining.

| **Antibody** | **Clone** | **Dilution or final conc.** | **Incubation time (min)** | **Supplier** | **Cat. No.** |
| --- | --- | --- | --- | --- | --- |
| IgM Fab AF405 |  | 10 µg/ml | 30 min on ice | Jackson (labelled in-house) | 115-007-020 |
| CD45R/B220 BV421 | RA3-6B2 | 1 µg/ml | 30 min at RT or o/n at 4 °C | BD | 562922 |
| CD45R/B220 PerCP/Cy5.5 | RA3-6B2 | 1 µg/ml | 30 min at RT or o/n at 4 °C | BioLegend | 103236 |
| MHC II (I-A/I-E) AF488 | M5/114.15.2 | 1 µg/ml | 30 | BD | 562352 |
| NFAT1 |  | 1:50 | 60 | Cell Signaling | 4389S |
| NF-κB (p65) | D14E12 | 1:400 | 60 | Cell Signaling | 8242S |
| p-BLNK (pY84) AF488 | J117-1278 | 1:20 | 60 | BD | 558444 |
| p-Myosin Light Chain 2 | T18/S19 | 1:200 | 60 | Cell Signaling | 3674S |
| p-Syk (Y525/526) | C87C1 | 1:50 | 30 | Cell Signaling | 2710S |
| Anti-mouse IgG (H+L) F(ab')<sub>2</sub> AF488 |  | 1:2000 | 60 | Cell Signaling | 4408S |
| Anti-rabbit IgG (H+L) F(ab')<sub>2</sub> AF488 |  | 1:2000 | 60 | Cell Signaling | 4412S |
| Anti-rabbit IgG (H+L) F(ab')<sub>2</sub> AF555 |  | 1:2000 | 60 | Cell Signaling | 4413S |
| Anti-rabbit IgG (H+L) F(ab')<sub>2</sub> AF647 |  | 1:2000 | 60 | Cell Signaling | 4414S |
| Phalloidin AF488 |  | 1:200 | 60 | Invitrogen | A12379 |
| DAPI ready-made solution |  | 1:1000 | 60 | Sigma-Aldrich | MBD0015 |

**Movie S1. Single-particle imaging of antigen-DNA complexes on DOPC.** Images were acquired with 10 ms exposure time and 0 s between frames.

**Movie S2. Single-particle imaging of antigen-DNA complexes on DPPC.** Images were acquired with 10 ms exposure time and 0 s between frames.

**Movie S3. Live-cell imaging of antigen extraction from bilayer substrates.** Dark spots appear when B cells rupture antigens from the DNA tether, separating an Atto647N fluorophore from a dark quencher. Extraction from DOPC is shown on the left, and from DPPC on the right. Images were acquired every 8 s. Scale bar: 5 µm.

